# Layer-specific ultra-high field 7T fMRI showing that threat anticipation is mediated by the pulvinar input to the superficial layers of primary visual cortex (V1)

**DOI:** 10.1101/431270

**Authors:** Ai Koizumi, Minye Zhan, Hiroshi Ban, Ikuhiro Kida, Federico de Martino, Beatrice de Gelder, Kaoru Amano

## Abstract

The perceptual system gives priority to threat-relevant signals with survival value. Its mechanism may not only include the processing initiated in the presence of threat signals but also in the mere anticipation of such signals. Here, we show that the pulvinar modulates activity in the early visual cortex (V1) specifically in threat anticipation. Using ultra-high-field 7T fMRI, we examined the layer-specific interaction between V1 and the pulvinar, while taking advantage of the fact that the directionality of such interaction is anatomically constrained in specific V1 layers. Only in anticipation of a fearful face target, but not of a control happy face target, was false perception of anticipated-yet-unpresented target face accompanied by stronger activity in the V1 superficial-cortical-depth (layers 1-3), which was preceded by pre-target-onset pulvinar activity. The pulvinar may contribute to the visual processing initiated in the anticipation of threat, and play an important role in anxiety.

## Introduction

Fear and anxiety are central issues in neuroscience, and they appear to partly emerge from the mechanisms supporting how the visual system uniquely processes sensory inputs with high behavioural relevance (1, 2) such as threat signals from facial or bodily expression (3–5).

The literature converges to postulate the pulvinar as a central relay, passing the threat-relevant sensory inputs forward for quick evaluation and behavioural response (6–9). In addition to the feedforward relay of sensory inputs, the pulvinar is also known to modulate activity in various cortical areas including the visual cortices (10–14). Such modulatory input from the pulvinar has been shown to be particularly critical in maintaining the responsiveness of the primary visual cortex, V1, to external inputs (11). However, whether and how the pulvinar-to-V1 input uniquely contributes to the visual perception of threat-relevant signals remains largely unknown. Whereas fear may be directly related to the actual presence of threat, threat anticipation is also critical for survival and is possibly related to sustained anxiety (15, 16). We speculated that the pulvinar-to-V1 modulatory input may play a role here.

We here tested the hypothesis that, the pulvinar-to-V1 input may help implement anticipation of threat-relevant signals into visual cortical processing. Our experiments used a simple task to detect a fearful face target (**Fig 1**), which serves as a social signal of threat (3–5). A fearful face detection task and a happy face detection task were conducted in separate sessions to establish a task context to anticipate either a fearful or happy face target in some, but not all, trials. In each task, neutral faces, instead of target emotional faces (fearful or happy), were presented on half of the trials. On those trials, participants falsely perceived either a fearful or happy face target at the rate of approximately 25%, depending on which facial emotion was anticipated in a given task. Unlike the percept in HIT trials where participants correctly perceived the presented fearful faces, the percept in false alarm (FA) trials could not be explained by sensory inputs of target faces but was instead likely due to anticipatory top-down processing.

**Fig 1.**
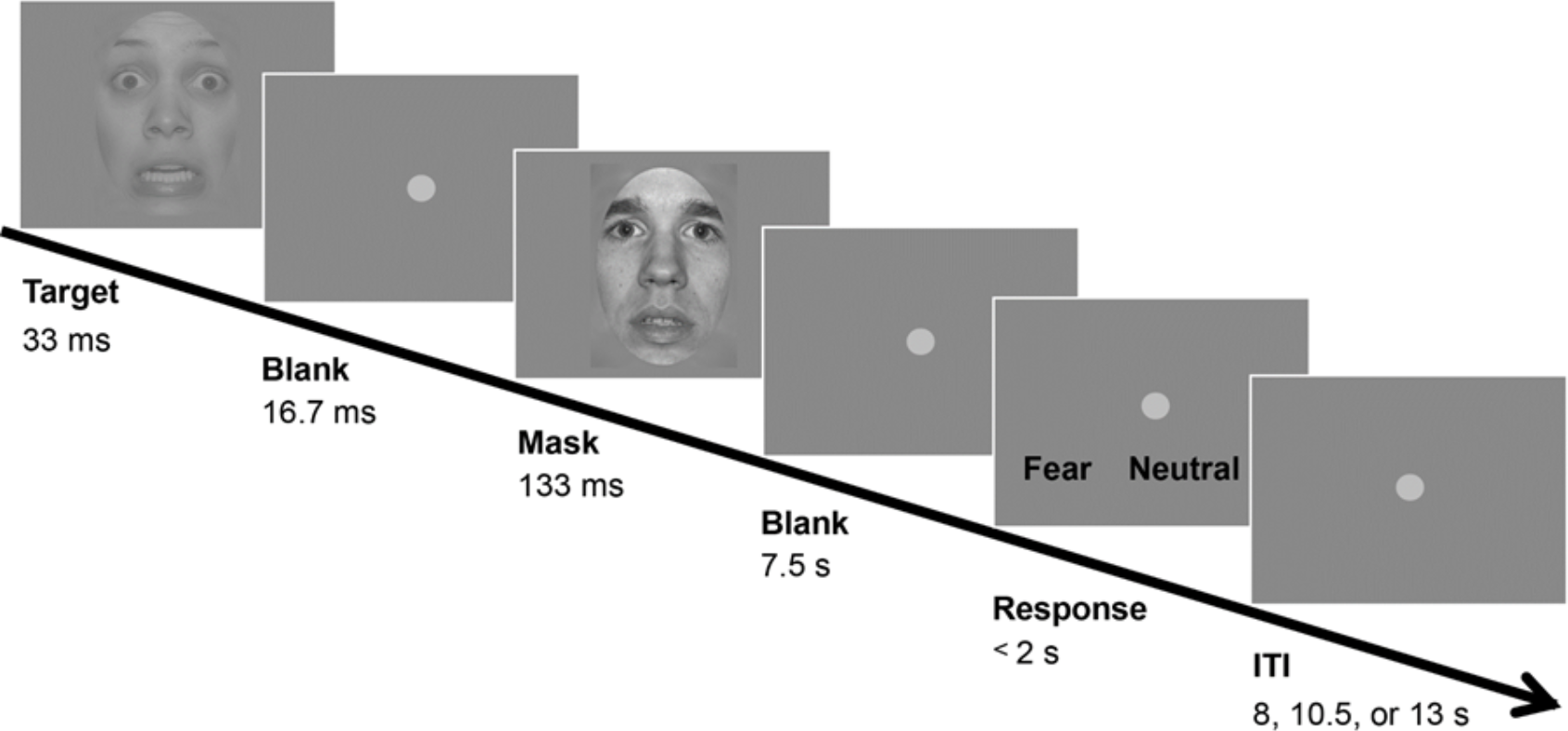
Design of the fearful face detection task. In each trial, either a fearful face target or a neutral face was presented briefly, followed by a mask. The mask was a neutral face with a different identity to the target face. Participants responded whether they perceived a fearful target or neutral face by pressing the response key, which was randomly assigned trial-wise. The control task used happy face targets and neutral faces (not shown), and otherwise identical procedures. ITI: inter-trial interval.

Using ultra-high-field 7T fMRI with spatial resolution of 0.8 mm, we explicitly examined the pulvinar-to-V1 input during FA trials while taking advantage of the anatomical constraint that the pulvinar sends modulatory input to V1 mainly in its superficial layers (13, 17). We found that FA of a fearful face, relative to that of a non-threatening happy face, was associated with stronger activity in V1 superficial-cortical-depth (roughly related to layers 1-3) (**Figs 2**, **S1, and S2**), even though the sensory inputs (neutral faces) were identical. Moreover, such V1 activity during the false percept of fearful face even numerically exceeded that during the correct percept of a fearful face, despite the fact that the actual sensory input of fearful face was only present in the latter.

The cross-correlation analysis showed that V1 activity in its superficial-cortical-depth during the false percept of fearful face, but not that during the false percept of happy face, was preceded by anticipatory pulvinar activity (**Figs 3** **and S3**). This pulvinar activity was already present prior to target face onsets (**Fig S4**), hence, could not be accounted for by the feedforward processing of and/or attention towards the presented faces. We observed no differential activity or cross correlation patterns related with the perception of fearful faces in a first-order thalamic area, lateral geniculate nucleus (LGN) as well as in a higher visual area (V4) (**Fig 4**), demonstrating some degree of specificity of our findings to the pulvinar and V1. Thus, our results suggest that the pulvinar sends input to the V1 superficial layers, which may bias the visual cortical processing towards perception of anticipated threat-relevant signals. This modulatory mechanism by the pulvinar may help elucidate how visual images of threat-relevant stimuli could emerge without corresponding sensory inputs, especially during flashback experiences of post-traumatic stress disorder (PTSD).

## Results

### Behavioural task

In each trial of the fearful face detection task, either a fearful face target or neutral face was briefly presented, followed by a neutral face mask (**Fig 1**). Participants then responded whether they perceived a fearful face target or a neutral face. A control task was performed in a counterbalanced order with detection of another emotionally salient but non-threatening happy face target. Perception of a fearful face reported on presentation of a fearful face target was classified as HIT, whereas perception of a fearful face reported on presentation of a non-fearful neutral face was classified as a false alarm (FA). The same designations were used for the happy vs. neutral faces in the control task.

### Measurement of cortical-depth dependent V1 activity and pulvinar activity with 7T fMRI

To non-invasively demonstrate pulvinar-to-V1 input during the task performance, we took advantage of the fact that such directionality is anatomically constrained in specific layers of V1. That is, while input from the pulvinar is sent to the superficial layers of V1, the deep layers of V1 send input to the pulvinar (13, 17). Specifically, we used ultra-high-field 7-Tesla fMRI with high spatial resolution (0.8 mm) to measure cortical-depth-dependent V1 activity while participants (N = 12) performed the detection tasks. Data for 1 participant were excluded from the analyses due to excessive head motion, and the data for the remaining 11 participants were analysed.

V1 activity was measured at 0.8 × 0.8 × 0.8 mm^3^ resolution and the data were preprocessed as previously described (18) to estimate the cortical depth dependent V1 activity (18–21) (Materials and Methods). We estimated activity in the superficial and deep cortical depths (**Figs 2A, 2B** **and 2C**), which roughly corresponded to cortical layers 1 to 3 and layers 5 to 6, respectively, due to the difference in thickness of each layer (22, 23). As the direct interactions between the pulvinar and V1 emerge primarily in V1 superficial and deep layers while avoiding the middle layer 4 (13, 17, 24), activity in middle depth was disregarded in the main analyses.

### Cortical-depth dependent V1 activity during fearful face perception

Our primary interest was how V1 activity in HIT and FA trials may differ between fearful and happy face detection tasks. We observed that whether higher activity was observed for the HIT or FA trials depended on the facial emotion anticipated in the tasks (fearful or happy) (**Fig 2D**). A repeated analysis of variance (ANOVA) revealed a second-order interaction between percept type (HIT/FA), emotion (fearful/happy), and cortical depth (superficial/deep) (*F*(1,10) = 5.667, *p* = 0.039). This interaction was due to the fact that percept type and emotion interactively modulated V1 activity only at superficial cortical depths (*F*(1, 10) = 7.840, *p* = .019), but not at deep cortical depths (see **Fig S1** for the deconvolved V1 time course and similar estimates of V1 peak activity with a different temporal window and **Fig S2** for the results separately plotted for individual participants).

At the superficial depth, V1 activity in FA trials of a happy face was less than that in HIT trials of a happy face (*t*(10) = 2.387, *p* = 0.038, *d* = 0.9). This result is consistent with a previous study demonstrating that V1 activity levels are typically greater for hit than for false alarm trials in a visual detection task with non-threatening targets (25), although cortical depth dependent activities were not reported.

Contrary to the happy face detection task, V1 superficial depth activity did not statistically differ between FA and HIT trials during the fearful face detection task (*t*(10) = −1.714, *p* = 0.117, n.s.). The trend that activity on FA trials was numerically larger relative to HIT trials may be because FA trials were accompanied by anticipation-driven activity in addition to activity driven by non-salient neutral faces, which in total became large enough to numerically exceed activity driven by inputs from the salient fearful face target in V1 level.

Interestingly, V1 superficial depth activity in FA trials of a fearful face was significantly greater than that in FA trials of a happy face (*t*(10) = −2.486, *p* = 0.032, *d* = 0.8), even though the same neutral faces were presented with the only difference being the task context to anticipate either a fearful or happy face target. This result suggests that V1 superficial depth activity may reflect excessive top-down processing in anticipation of fearful face targets. At deep cortical depths, V1 activity was not modulated by percept type or facial emotion.

**Fig 2.**
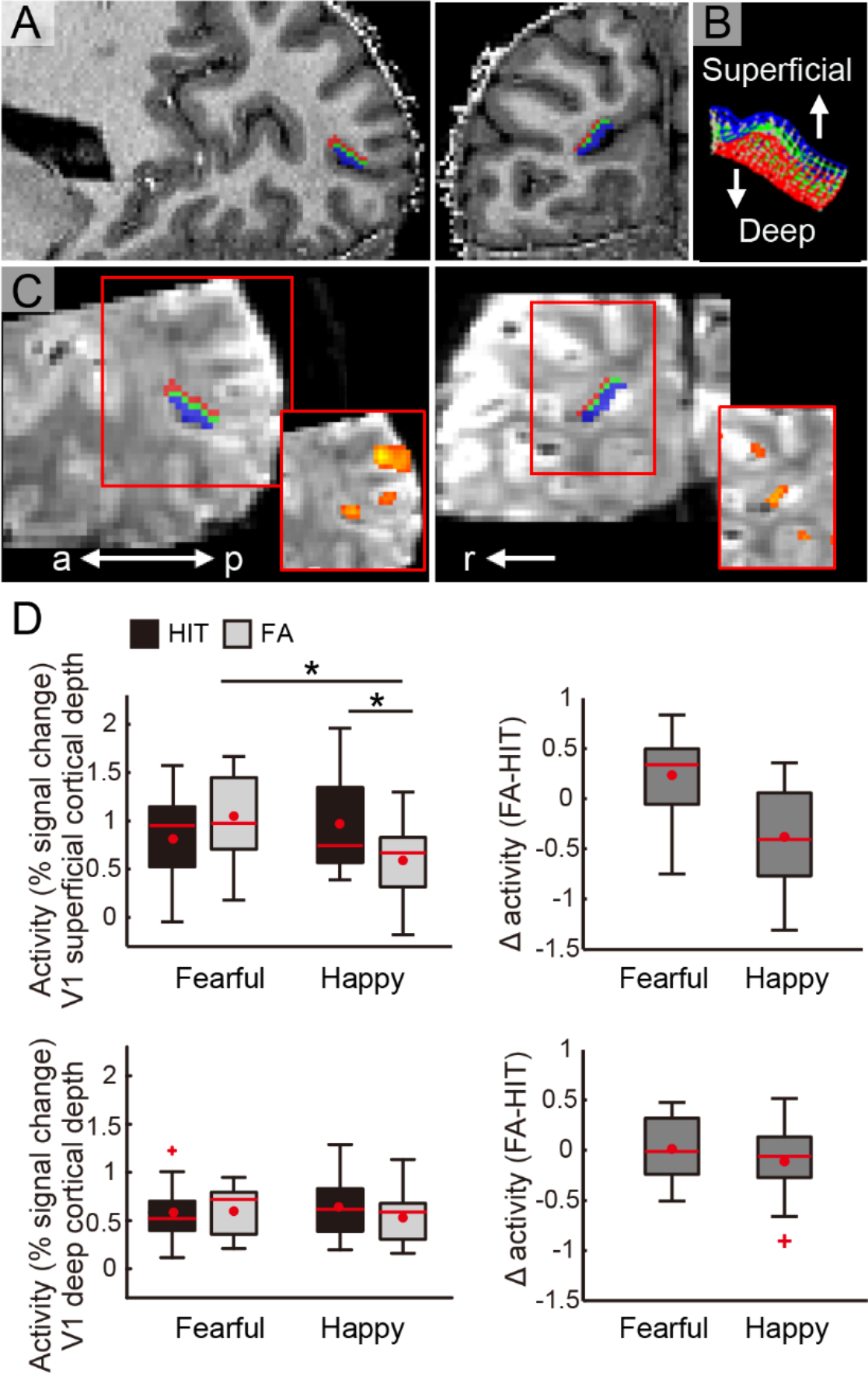
Demonstrations of V1 cortical depths and cortical depth-dependent activity during the tasks. **A.** V1 cortical depth is visualised on an anatomical image of a representative participant with sagittal (left panel) and coronal views (right panel). The voxels allocated to superficial (outwards to pial surface) and deep (inwards to white matter) cortical depths are shown in blue and red, respectively. The voxels allocated to the intermediate depth (shown in green) were disregarded in the main analyses (see *Materials and Methods*). **B.** The cortical grid mesh within which the voxels were allocated. **C.** The V1 cortical depth is visualised on 3D EPI images in sagittal and coronal views (left and right, respectively). Red squares at the lower right demonstrate activity for all face targets relative to baseline on the EPI smoothed with a 3D kernel of 2.4 mm full width at half maximum (FWHM). **D.** Peak activity at V1 superficial and deep cortical depths (upper and lower rows, respectively) in HIT and false alarm (FA) trials in the fearful face and happy face detection tasks. The difference in activity between FA and HIT trials are demonstrated in the right panel for each task, with a larger value indicating greater activity for FA than for HIT trials. Box plot shows upper (75%) and lower (25%) quartiles with median (red line) and mean (red dot), with whisker showing maximum and minimum value. An outlier (outside of ± 2.7 standard deviations within a distribution for a given condition) is shown with a red cross. a: anterior, p: posterior, r: right, \**p* < 0.05. See also **Fig S1** for the deconvolved time course and estimates of peak activity in V1 with a different temporal window and **Fig S2** for the activity levels separately plotted for each participant.

### Pulvinar-to-V1 input during fearful face perception

We next examined whether enhanced V1 activity for FA trials of fearful faces was related to inputs from a higher-order thalamic area, the pulvinar (**Fig 3A**). The pulvinar has been suggested to contribute to the coarse feedforward pathway (“low load”) for threat perception (3, 6, 8). However, the pulvinar also receives various inputs from higher cortex including prefrontal areas (10, 26), and may play a role in higher-level perceptual processing (12, 27, 28). Thus, we predicted that pulvinar input may act to modulate V1 activity in FA trials of fearful faces. Here, the pulvinar was defined as the area within the anatomical boundaries (29–31) that responded to the onsets of faces on all trials, i.e., unbiased towards a certain percept type and/or facial emotion (see *Materials and Methods* for details).

We observed that pulvinar activity was greater on FA trials than on HIT trials (**Fig 3B**), as demonstrated by a main effect of percept type in a repeated measures ANOVA (*F*(1,10) = 8.310, *p* = 0.016). However, this activity difference was not specific to FA trials of fearful faces, as there was no significant interaction between percept type and emotion (*F*(1,10) = 0.155, *p* = 0.702). The pulvinar may show enhanced activity with a false percept in general, as has been reported in a previous study during a false detection of change in non-emotional stimuli (32) (see also **Fig S3** for the deconvolved time course of the pulvinar and estimates of pulvinar peak activity with a different temporal window). The enhanced pulvinar activity on FA trials could not be explained away by mere error signals in perception, as a control analysis showed that pulvinar activity was not significantly enhanced on errored Miss trials relative to error-free CR trials (**Fig S5**).

More importantly, while pulvinar activity alone did not dissociate between fearful and happy faces (**Fig 3**), a time-shifted cross correlation analysis (see *Materials and Methods*) suggested that there was enhanced input from the pulvinar to V1 specifically in FA trials relative to HIT trials of fearful faces. We conducted this correlational analysis based on the assumption that a direction of interaction between two brain areas can be inferred when activity in one brain area shows stronger correlation with activity in another area from a certain past time point (33). Our results suggested that there was an enhanced pulvinar-to-V1 correlation in FA trials of fearful faces with a time lag of approximately 1500 ms, and that this was significantly stronger than its counterpart in HIT trials of fearful faces (*t*(10) = 4.053, *p* = 0.018, and *t*(10) = 3.541, *p* = 0.045 when pulvinar activity precedes by 1875 ms and 1250 ms, respectively, Bonferroni corrected) (**Fig 3C**). The suggested directionality of the interaction is consistent with the known anatomical constraint that the pulvinar sends inputs to V1 predominantly in its surface layers (13, 17).

Considering that the observed time lag (i.e., 1500 ms) was relatively long, it is unlikely that the interaction reflected direct information transmission between the two areas, because input from pulvinar appears to reach V1 within a few hundreds of milliseconds (11). Rather, such a long time lag is likely to reflect the indirect interaction in which pulvinar potentiated the responsiveness of V1 in its superficial depth, which later resulted in the enhanced activity in V1 when the sensory inputs (i.e., neutral faces) were presented.

Supporting such possibility, analysis of activity prior to target face onset revealed that pulvinar activity was already enhanced prior to FA trial of a fearful face relative to that of a happy face (**Fig S4**), resembling the pattern of activity differences in V1 superficial cortical depth which was observed following target face onset (**Fig 2D**). Unlike the pulvinar, there was not yet any differential activity in V1 superficial cortical depth during the pre-onset period, further supporting the view that anticipatory pulvinar activity preceded enhanced activity in V1 superficial cortical depth towards the presented neutral faces, potentially contributing to false percept of fearful faces. Such pulvinar-to-V1 input may play a selective role in top-down modulation of visual processing towards perception of fearful face targets, as there was no significant difference in correlation between HIT and FA trials of happy faces (**Fig 3C**).

It has been previously shown that the pulvinar has subdivisions that are distinctively connected with different cortical areas. Specifically, whereas the lateral pulvinar has direct connections with visual cortices including V1, the medial pulvinar has direct connections with prefrontal areas (10). Given this, we additionally examined whether there was differential activity across the subregions of the pulvinar, which were anatomically defined with a histological atlas (34). The results suggested that one lateral and one medial subregion showed significantly larger activity on FA trials of a fearful face relative to FA trials of a happy face (**Fig S6**), similarly to V1 activity at the superficial cortical depth (**Fig 2D**). These results hint at the possibility that task-driven anticipation of threat signals may be coded in the medial pulvinar through its interaction with the prefrontal areas where task-set is generally represented (35), while the lateral pulvinar may directly modulate V1 based on anticipation. However, as the results within these subregions were limited in that there was no significant interaction between conditions (HIT/FA) and emotions (fearful/happy), these results should be treated as indicative.

**Fig 3.**
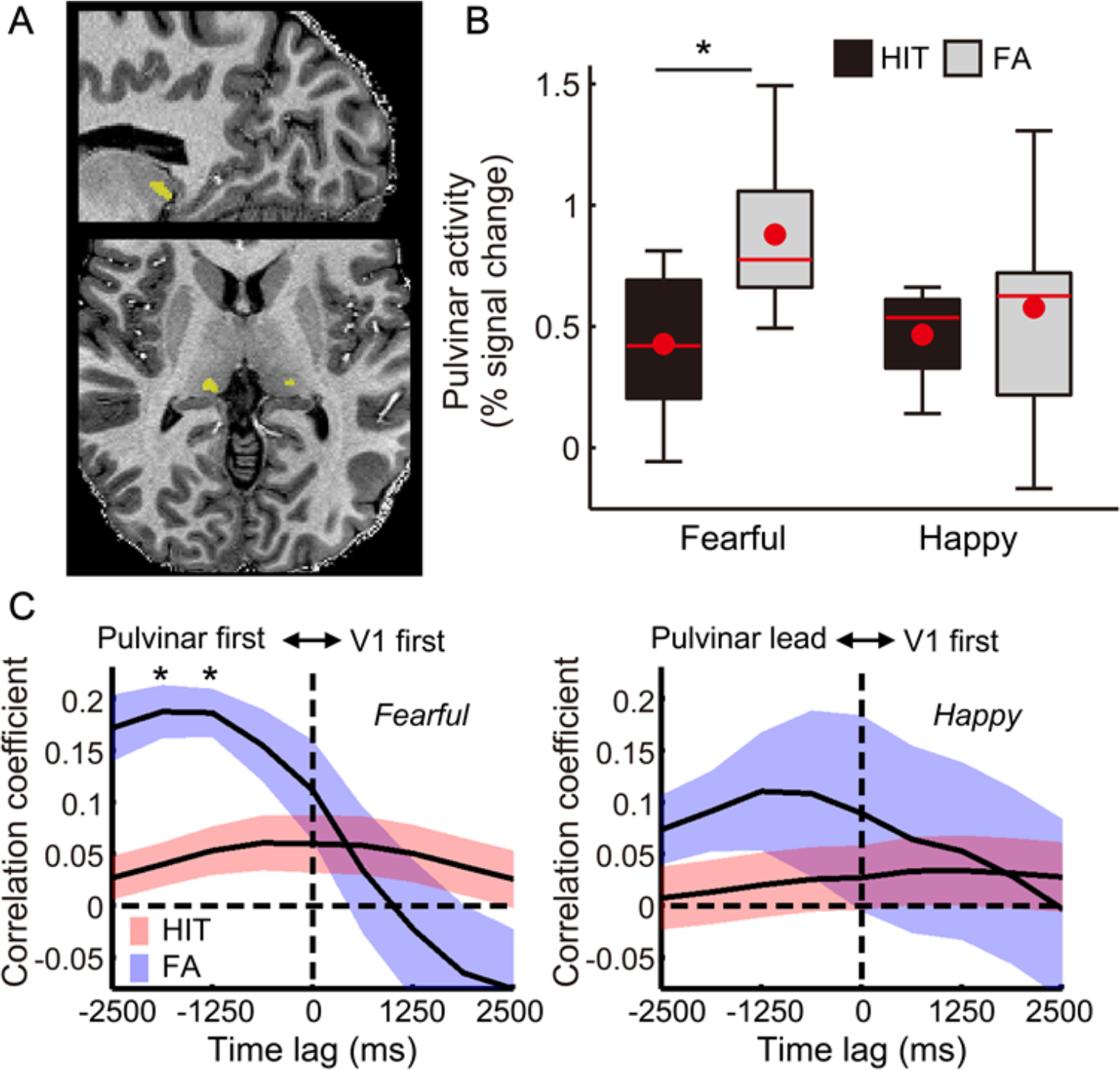
The pulvinar showed enhanced activity, which preceded V1 superficial cortical depth activity on false alarm (FA) trials of fearful faces. **A.** Demonstrations of pulvinar ROIs from a representative participant (see *Materials and Methods* for ROI definition). **B.** Pulvinar showed enhanced activity in FA trials relative to HIT trials during the fearful face detection task (*t*(9) = −2.704, *p* = 0.024, *d* = 0.7) but not during the happy face detection task (*t*(9) = −0.649, *p* = 0.533, n.s.). There was no significant difference between FA trials of the fearful face detection task and that of the happy face detection tasks (*t*(9) = 1.632, *p* = .137, n.s.). There was no significant interaction between percept type and emotion (*F*(1,9) = 1.484, *p* = 0.254). Here, data from one participant with an extreme outlier (activity exceeding 5 standard deviations above the group mean across four conditions, i.e., HIT/FA x fearful/happy) were excluded. When this participant’s data were included in the analysis, the statistical results remained qualitatively similar (see **Fig S3C**). Box plot shows upper (75%) and lower (25%) quartiles with median (red line) and mean (red dot), with whisker showing maximum and minimum value. **C.** The time-shifted cross correlation analysis suggested that the pulvinar preceded V1 superficial cortical depth activity in FA trials of the fearful face detection task (left panel). There was no significant difference in correlation between HIT and FA trials of the happy face detection task (right panel). Error bar and color shaded area indicate standard error of mean. \**p* < 0.05. See also **Figs S3-8**.

### Control analyses

To examine the specificity of the aforementioned results, we conducted three control analyses. First, we examined the potential contributions of higher visual cortex, V4. Here, V4 was defined similarly to V1 along with the anatomical landmarks (36) (see **Fig 4A** and *Materials and Methods*). We selected V4 as a control area because other higher visual areas such as fusiform areas were outside of the fMRI coverage and we had difficulty locating task-relevant voxels within other earlier visual areas such as V2/3 in a similar manner as V1. V4 it is known to have a direct connection with V1 (37) including its input to the V1 superficial layers (38, 39), and may thus additionally contribute to V1 superficial depth activity in FA trials of fearful faces.

Contrary to this possibility, V4 did not show any differential activity (**Fig 4B**) or cross correlation with V1 activity across the percept types (**Fig S7C**). Note that these null results do not exclude the possibility that other areas besides the pulvinar also modulated V1 activity during FA trials of fearful faces. Future studies should investigate whether and how pulvinar-driven V1 activity interacts with activity in other higher areas, such as the fusiform face area and prefrontal areas. Although whole brain analyses may help address such questions, analyses were not performed because of the restricted brain coverage (see *fMRI Data acquisition* in *Materials and Methods*), and because the spatial distortion and/or signal-to-noise ratio (SNR) were expected to be highly inhomogeneous even across the covered brain areas due to ultra-high magnetic field.

As a second control analysis, we examined activity of a first-order thalamic region, the LGN (**Fig 4C**), which was localised in the same manner as the pulvinar (see *Materials and Methods*). Similarly to the pulvinar, the LGN is known to have bidirectional connections with V1 (13, 17, 24). However, because these connections do not include the V1 superficial layers, superficial depth activity in FA trials of fearful faces is unlikely to be related to the LGN. Consistent with this idea, our analysis showed that neither the activity level of the LGN (**Fig 4D**) nor the time-shifted cross correlation between the LGN and V1 were altered in FA trials of fearful faces (**Fig S7B**).

As a third control analysis, we examined whether similar results to those observed in FA trials of fearful faces would be present in MISS trials, in which a fearful face was presented but not detected. While the aforementioned results for FA trials of fearful faces may reflect enhanced top-down processing as we speculated earlier, other non-mutually exclusive possibilities are worth considering. Specifically, the results in FA trials may reflect a mere mismatch between the sensory input and the reported percept, in which case similar results as FA trials would be expected in MISS trials. Contrary to this possibility, we did not observe any notable results specific to MISS trials of fearful faces either in V1 or in the pulvinar (**Fig S5**), excluding the possibility that the results in FA trials of fearful faces merely reflected a mismatch between the input and percept.

Taken together, the results of these control analyses suggest some degree of specificity of the pulvinar and V1 superficial cortical depth in their involvement in false perception of fearful faces.

**Fig 4.**
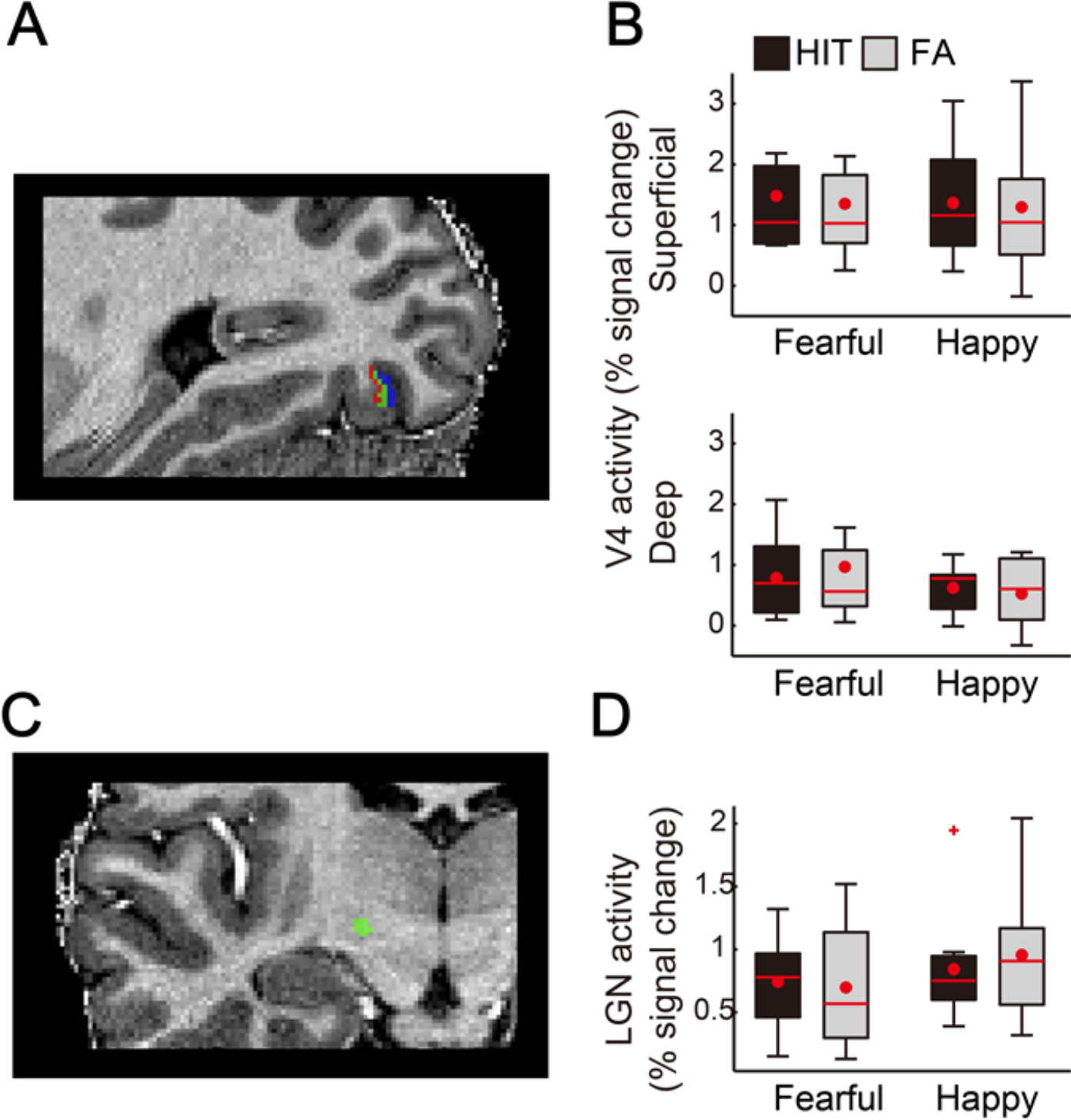
Control analyses showing no differential activity in V4 and the LGN in HIT compared with false alarm (FA) trials. **A.** V4 cortical depth is visualised on the anatomical image of an example participant, with the same colour coding as for V1 shown in **Fig 2**(see *Materials and Methods* for ROI definition). **B.** Unlike V1, V4 did not show any differential activity between the percept types and facial emotions, regardless of cortical depth. A second-order interaction between percept type, emotion, and cortical depth was non-significant (*F*(1,10) = 1.028, *p* = 0.335). The main effects of percept type and emotion were also non-significant (*F*(1,10) = 0.124, *p* = 0.732; *F*(1,10) = 0.850, *p* = 0.37, respectively), while only the main effect of cortical depth was significant (*F*(1,10) = 18.523, *p* = 0.002). **C.**Demonstrations of the LGN ROI from an example participant (see *Materials and Methods* for ROI definition). **D.** Unlike the pulvinar, the LGN showed no differential activity between the percept types and facial emotions. Neither the main effect of percept type nor that of emotion was significant in a repeated-measures ANOVA (*F*(1,10) = 0.100, *p* = 0.759; *F*(1,10) = 3.014, *p* = 0.113, respectively). The two-way interaction was also non-significant (*F*(1,10) = 0.363, *p* = 0.560). Box plot shows upper (75%) and lower (25%) quartiles with median (red line) and mean (red dot), with whisker showing maximum and minimum value. An outlier (outside of ± 2.7 standard deviations within a distribution for a given condition) is shown with a red cross. See also **Fig S7**.

## Discussion

In summary, we observed that the false percept of a fearful face relative to that of a happy face was accompanied by increased activity at superficial cortical depths of V1, which constitutes the earliest stage of the visual cortical hierarchy. Moreover, this V1 activity was likely to be driven by anticipatory activity in the pulvinar, consistent with the anatomical constraint that V1 receives modulatory input from the pulvinar predominantly in its superficial layers (13, 17). This mechanism by which the pulvinar modulates visual cortical processing may particularly contribute to the perception of threat signals, as it did not generalise to the perception of non-threatening signals, i.e., happy faces.

The current findings are not mutually exclusive with the traditional view emphasizing the role of the pulvinar in the subcortical route (“low load”) to process the presented threat signals in a feedforward manner. The subcortical route is thought to bypass the cortex to rapidly relay the retinal input to the amygdala via the superior colliculus and pulvinar (3, 6). Such coarse feedforward processing has been speculated to result in erroneous, false perception of threat signals (6). Although the role of such a subcortical route has been demonstrated in humans (9, 40, 41), its function may be relatively degraded in humans compared with other species such as rodents (8, 10). Our study suggests that, the pulvinar may play an additional role to modulate activity in the early stage of visual cortex in anticipation of threat-relevant signals in humans.

Thus, the pulvinar may play parallel roles in the processing initiated by the presented threat signals and that head started by prior anticipation of such signals, which are both likely to ensure the perception of threat-signals with survival values.

It may be counterintuitive to expect a crucial role of V1 in the perception of fearful faces, given that V1 is fine-tuned to low-level visual features that would constitute only subcomponents of fearful faces. Nevertheless, recent findings suggest that a lower visual cortical area can reflect higher-level features, when the prediction signals for such features originate from a higher area (42). While such top-down signals are typically expected to descend from higher cortical areas, it has recently been speculated that the pulvinar also contributes to such top-down signals (27), including contextual signals (12). Given that the pulvinar appears capable of coding threat signals, including a complex fearful face (3, 8), it may modulate early stage processing in V1 to immediately bias visual cortical processing towards threat perception.

How false perception of non-threatening cues, such as happy faces, emerges remains to be investigated in future studies. One possibility is that such percept would reflect top-down modulation of visual cortical areas higher than V1, similarly to the processing of facial identity (42). As higher visual areas were not covered in this study (to allow for the spatial resolution desired), future study may directly compare the mechanisms underlying false perception of threat-relevant cues versus non-threatening cues. Moreover, future studies may examine the potential involvement of other brain areas expected to be critical for threat perception, such as the amygdala, which were not covered here.

The current results, together with the known directionality of input from the pulvinar to V1 superficial layers (13, 17, 24), converge to support a critical role of pulvinar input to V1 in visual threat perception. Such modulatory input from the pulvinar may contribute to the perception of threat-relevant images without corresponding sensory inputs as in clinical cases of flashbacks reported in PTSD.

## Materials and Methods

### Participants

We enrolled 12 participants in this study (6 male, mean age 23.73 ± s.d. 3.64, 2 left-handed) after providing written consent. Participants were all healthy and had normal or corrected-to-normal vision. They completed 2 sessions, in which fearful and happy face targets were presented in a counterbalanced order and were paid 60 euro per session.

One participant was removed from analysis due to excessive head motion (mean across runs > 4 mm). Scanning was conducted at the Maastricht Brain Imaging Centre, Maastricht University, Netherlands. The protocol was approved by the ethical committee of Maastricht University.

We estimated that a sample size of N = 11 would be sufficient to detect a medium to large effect size (f = .30) with an alpha power of 0.05 and power of 80% (G*Power version 3.1.9.2). Although we initially aimed for N = 12 to be conservative, one participant was removed from analysis as described above, leaving us with N = 11.

### Stimuli

Face images of 6 models (3 male) displaying fearful, happy, and neutral expressions were taken from the NimStim face stimulus set (43). We only included face images with an open mouth so that the local feature of an opened mouth alone would not enable detection of a fearful or happy face rather than a neutral face. The images were grey-scaled and cropped into oval shapes to eliminate hair. They were then matched for luminosity, contrast, and spatial frequency spectrum with the SHINE toolbox (44) implemented in MATLAB (R2011b, The Mathworks, Natick, Massachusetts, USA). We refrained from further manipulation of stimulus properties, as excessive manipulation itself could unintentionally induce differential activity in V1, which is sensitive to lower-level stimulus properties.

### Procedure

The fearful face detection task required participants to detect a briefly presented fearful face followed by a mask (neutral face). On each trial, either a fearful or neutral face appeared as a target for 33 ms. Following a blank of 16.7 ms, a neutral face was presented for 133 ms as a mask that rendered the target face less visible but did not completely abolish its visibility. The model for the mask face was different from the model for the target face on every trial.

To further control task difficulty, the contrast of the target face was reduced to 35% of the contrast of the mask face, as determined during our pilot study. After 5 s had elapsed from the target onset, response key assignment was shown on the screen. Responses for a fearful or neutral face were assigned to either the left or right key, which was randomized across trials. Participants were instructed to respond with their first guess whether they had perceived a fearful or neutral face target, and to quickly respond with their right hand within a 2 s time window. After a randomized inter-trial-interval from the onset of response assignment (8, 10.5, or 13 s), another trial was initiated. There were 24 trials in each of 8 runs (8 min 10 s per run), comprising 12 trials each for fearful and neutral face targets. The order of trials was randomized.

The control task with happy faces was conducted in an identical manner during another scanning session, except that the fearful face targets were replaced by happy face targets using the same models. The neutral face images remained identical. The fearful and happy face sessions were conducted on two separate days in a counterbalanced order across participants.

Stimuli were presented with Psychtoolbox (45) implemented in Matlab (R2012a, the Mathworks). A separate localizer run (3 min 30 s) was acquired to define V1 (see *V1 localization*).

### Data analyses

We conducted subject-level analyses of fMRI data in BrainVoyager 20.2 (Brain Innovation, Maastricht, the Netherlands) (see *fMRI processing*), and subsequently, conducted group-level repeated measures ANOVAs in IBM SPSS Statistics (version 18). The assumption of sphericity was met in the current dataset because there were only two levels per factor (e.g., emotion) in each repeated ANOVA.

Following previous studies (46, 47), when an omnibus 3-way repeated measures ANOVA revealed a significant 3-way interaction, we conducted 2-way (n-1 way) repeated measures ANOVAs to locate a simple interaction effect. A significant 2-way interaction that was identified was subsequently tested with a t-test (two-tailed) to examine simple main effects. Similarly, when an omnibus repeated measures ANOVA initially involved only two factors (i.e., 2-way), a significant interaction was similarly followed by t-tests (two-tailed). The series of t-tests were not susceptible to the inflation of type 1 error as they followed significant interactions in the initial omnibus ANOVAs, as has been validated and commonly practiced previously (46, 48).

We estimated effect sizes in the post-hoc t-tests with Cohen’s d, where a value larger than 0.8 indicated a large effect size and a value larger than 0.5 indicated a medium effect size (47, 48). All t-tests reported in this manuscript were two-tailed.

### Behavioural performance

Behavioural performance was also analysed in IBM SPSS. The ratio of FA was similar between the fearful face detection task and the happy face detection task (M = 28.51 ± 3.52%; M = 24.13 ± 3.07%, respectively), and the rates in each task did not significantly differ from each other (*t*(10) = 1.845, *p* = 0.095). The ratio of HIT trials was higher for the happy face detection task (M = 60.36 ± s.e. 4.36%) than for the fearful face detection task (M = 49.18 ± 4.42%) (*t*(10) = - 4.155, *p* < 0.01). This is consistent with previous literature showing that explicit labelling of happy faces is easier than that of negative faces (49). As described in *Stimuli*, we avoided any manipulation to equate perceptual performance between the tasks, such as morphing of facial images (4), to prioritize the measurement of V1 activity, which is sensitive to changes in lower-level stimulus properties.

As discussed in *Introduction*, we speculated that FA may emerge when top-down anticipatory processing amplifies the percept to compensate for the absence of sensory inputs. Although anticipatory processing may also contribute to the percept in HIT trials, such a contribution is expected to be smaller than that of FA trials given that at least some portion of HIT trials are attributed to the presence of sensory inputs. This can be speculated when simply considering the proportion of HIT trials that are mainly driven by sensory inputs or alternatively by anticipation. That is, while the proportion of HIT trials (≒ 55%) that is the same as the total FA rate (≒ 25%) may be attributed to amplified anticipatory processing, the remaining HIT trials (≒ 30% out of 55%) is likely to reflect sensory inputs for the target faces instead. Thus, FA trials are more likely to reflect top-down related processing than are HIT trials on average.

### fMRI data acquisition

MRI data were acquired with a 7T Magnetom scanner (Siemens, Erlangen, Germany) at the Scannexus facility located at the Department of Cognitive Neuroscience, Faculty of Psychology and Neurosciences, Maastricht University (NL), with a Nova 32-channel receive head coil (Nova Medical, Wilmington, USA). For functional data acquisition, 2D gradient-echo planar images (EPI) were acquired at 0.8 mm isotropic resolution, with the following parameters: repetition time (TR) = 2500 ms, echo time (TE) = 21.8 ms, flip angle = 80, GRAPPA acceleration factor = 3, matrix size = 154 × 236, field of view (FOV) = 123 mm × 188 mm, slice thickness = 0.8 mm, number of slices = 40, no gaps. The slices were oriented to cover both the pulvinar and V1. To achieve maximal brain coverage with these parameters, right to left (RL) phase encoding was used for the task runs, so that the temporal areas outside the FOV were folded within the FOV to be trimmed later offline. A run of 5 TRs with the same parameters but with the opposing left to right (LR) phase encoding direction was acquired immediately before each task run for top-up EPI distortion correction of the functional images during offline analysis (see *fMRI processing* for more details).

For anatomical data in 9 participants, a T1-weighted scan and a proton density-weighted scan were acquired with a resolution of 0.6 mm isotropic (FOV = 229 mm × 229 mm, matrix size = 384 × 384, flip angle = 5. T1-weighted: TR = 3100 ms, TE = 2.52 ms; proton-density-weighted: TR = 1440 ms, TE = 2.52 ms). For the other 3 participants, we used anatomical images with a spatial resolution of 0.7 mm from previous unrelated experiments.

### fMRI processing

fMRI analyses were conducted in BrainVoyager 20.2 (Brain Innovation, Maastricht, the Netherlands). For preprocessing of fMRI data, we trimmed the lateral sides of EPI images by a small amount (60 voxels) to remove the folded-in tissue outside the FOV. The folding-in and trimming did not affect the coverage of the bilateral pulvinar and V1. The trimmed EPI images were then slice time corrected (sinc interpolation) and corrected for 3D rigid body motion (trilinear/sinc interpolation). Distortions of the EPI images from the task runs were adjusted against EPI images taken immediately before each task run with the opposing encoding phase, with the BrainVoyager plugin COPE (50) (http://support.brainvoyager.com/available-tools/49-available-plugins/477-cope-plugin-for-epi-distortion-correction.html).

EPI images then underwent temporal high-pass filtering with 2 cycles. EPI images were manually aligned to the anatomical images in BrainVoyager, while optimising alignment around V1 and the pulvinar. Among the 8 runs for each of the main tasks with fearful and happy face targets, EPI images from the runs with 3D motion larger than 2.5 mm were discarded from analyses, because large motion induced excessive and/or unique EPI distortion that interfered with precise alignment and the subsequent cortical depth specific analyses. One participant’s data were excluded from further analysis due to excessive head movements (> 4 mm). For the remaining participants, the run numbers included in analyses did not differ between the fearful face task (M = 7.64 ± s.d. 0.67) and the happy face task (M = 7.45 ± 0.82) (*t*(10) = 0.482, *p* = 0.640).

Task-related activity was then estimated with a deconvolution analysis, in which responses for successive 5 points (2.5 s × 5 TRs) were estimated, starting from the onset of the target face for each of the 8 experimental conditions (fearful face hit, miss, false alarm, correct rejection, as well as happy face hit, miss, false alarm, and correct rejection). The temporal peak of the deconvolved time course at 5 s (2 TRs) from the target face onset was used in the analyses described in *Results*. The entire deconvolved time course and estimates of peak activity with a different temporal window in V1 and the pulvinar are shown in **Figs S1 and S3**. To remove the influence of task-response-related activity from target face related activity of interest, the model also included the estimates for the 5 data points starting from the onset of the response period for each of the 8 conditions. Additionally, to remove the influence of head motion, the model included an additional 6 nuisance parameters in 3 translation directions and 3 rotation axes. EPI were not spatially smoothed to maintain laminar specificity.

### V1 localization

To localize V1 in each participant, a retinotopy scan was acquired. During the scan, colour/luminance-flickering wedge-shaped checkerboard patterns (30 deg in polar angle) were presented along the horizontal or vertical meridian alternately for 15 s each with 6 repetitions, as per the previously validated procedure (51). Checkerboard patterns were flickered at 4 Hz and were displayed in 1 of 4 colour combinations (red/green, blue/yellow, black/white, and magenta/cyan) to activate neurons with various response profiles to enhance the signals to identify V1 boundaries. The boundary of V1 was delineated with a general linear model comparing activity between the horizontal and vertical presentation periods as previously described (51).

### Anatomical image processing and cortical depth specific estimation of V1 activity

Inhomogeneity of T1-weighted images was corrected by dividing the original image intensities by the proton density images (52). Subsequently, the corrected T1 image was resampled at a resolution of 0.8 mm to match the resolution of EPI. The boundaries of grey-white matter and the pial surface were first estimated with BrainVoyager 20.2, and further corrected manually to improve the precision and to remove the blood vessels and dura mater based on image intensity. The anatomical image was not transformed to standardized coordinates but was kept in native space to maintain its laminar properties undistorted.

In order to specifically select task-relevant voxels within the retinotopically localised V1 (but not for subsequent estimation of cortical depth dependent activity), functional images from the task runs were smoothed with a gaussian kernel of 2.4 mm full width at half maximum (FWHM) to avoid potential bias in selecting voxels based on the activity at one cortical depth over the others. Note that task-relevant voxels were selected based on the target face onsets from all trials including all conditions, and therefore were not biased towards one particular condition over another. The task-relevant voxels were selected within the localised V1 of each hemisphere. Specifically, the task-relevant voxels were defined based on the contrast between all target face onsets versus baseline at a threshold of *p* < 0.001 uncorrected, while considering a haemodynamic delay of 5 s (2 TRs × 2.5 s). The task-relevant voxels were successfully located in both hemispheres in 8 participants, while they were located in only one hemisphere in the remaining 3 participants (right only, N = 1; left only, N = 2). For participants with peaks located in both hemispheres, the estimates of V1 activity were averaged between the hemispheres.

To define the cortical depths, we used the Laplace equation to estimate cortical thickness and then obtained an equidistant definition of depth with respect to the local thickness (18, 53) at three depth levels (from 25, 50, and 75% depth levels relative to the local cortical depth in an inward direction) centering around the spatial activity peak (i.e., voxel with highest activation level) among the pre-defined task-relevant voxels with 15 × 15 grids of 0.5 voxels. This covered an average of 22.67% (s.e. = 3.95) of the retinotopically localised V1 (see *V1 localization*). Individual voxels were assigned to the adjacent cortical depth, and were used as regions of interest (ROIs) in the subsequent analyses (**Fig 2**). We confined our analyses to voxels allocated to the superficial and deep cortical depth groups, which roughly correspond to cortical layers 1 to 3 and layers 5 to 6, respectively, due to the difference in anatomical thickness of each layer (22, 23). The activity in the middle depth was disregarded in the main analyses, given that the primary direct interactions between the pulvinar and V1 take place in the V1 superficial and deep layers while avoiding the middle layer 4 (13, 17, 24).

One potential limitation of the current method is that the gradient-echo imaging sequence employed here is known to have a better SNR towards the surface of the cortex (53). This SNR difference may mean that the study was underpowered to elucidate potential additional contributions of V1 at the deep cortical depth. However, this cannot account for the main result that the percept type and the facial emotion interactively modulated V1 activity at its superficial depths (**Fig 2D**). Moreover, a previous study with similar 7T fMRI parameters has successfully elucidated significant effects in V1 even at its deep cortical depths (23). In the current study, we were also able to detect a significant difference between some experimental conditions (i.e., HIT and CR of a happy face) in the V1 deep cortical depth (**Fig S8**), suggesting that there was sufficient power to detect some difference in activity even at the deep cortical depth albeit the potential SNR difference. As mentioned in *Discussion*, the current results, together with the known directionality of the interaction from the pulvinar to V1 superficial layers (13, 17, 24), support a role of the pulvinar to guide V1 activity in visual threat perception.

### Localization of the pulvinar, LGN, and V4

In addition to the aforementioned V1, we localised the pulvinar, as well as the LGN and V4 as control regions. Each of the thalamic regions (the pulvinar and LGN) was first anatomically defined based on their physical properties (29–31) in the high-resolution T1 resampled at 0.8 mm in native space. Task-relevant voxels within each thalamic region were then defined in a similar manner as for V1, based on activation at the temporal peak of the time course (5 s from the target face target onset) that was larger than baseline at a liberal threshold of *p* < 0.01 to compensate for a generally lower SNR in subcortical areas.

Pulvinar ROIs were located in both hemispheres for all participants except for 1 participant with a ROI in the left hemisphere only. LGN ROIs were located in both hemispheres for 5 participants, while they were located in either the right or left hemisphere for 4 and 2 participants, respectively. For participants with pulvinar and/or LGN ROI(s) located in both hemispheres, estimates of activity were averaged between the hemispheres. The ROI for V4 was localised based on data from the retinotopy scan in a similar manner as for V1 localization (see *V1 localization*), while additionally referring to the predefined anatomical landmarks of V4 (36). V4 ROIs were located in both hemispheres for all participants. All ROIs were defined with EPIs spatially smoothed with a gaussian kernel of 2.4 mm FWHM to enhance SNR.

Additionally, ROIs for the pulvinar subregions were defined with a previously reported histological atlas (34). The atlas was imported to BrainVoyager, and the entire pulvinar including all subregions was manually aligned to the pulvinar in each participant’s Native space. We included the subregions that compose the lateral, inferior, and medial portions of pulvinar which are widely implicated in visual processing (8, 10), namely the nucleus pulvinaris oromedialis (lateral), nucleus pulvinaris orolateralis (lateral), pulvinar laterale (lateral), nucleus pulvinaris intergeniculatus (inferior), nucleus pulvinaris (medial), and pulvinar mediale (medial). The entire voxels within each subregion were used to estimate the activity level shown in **Fig S6**.

We performed ROI-based analyses instead of whole brain analyses because the coverage of the brain was limited but this allowed for higher spatial resolution (see FOV in *fMRI data acquisition*). Moreover, spatial distortion and/or the SNR were expected to be inhomogeneous even across the covered brain areas due to ultra-high magnetic field.

### Analyses of pre-onset activity and time-shifted cross correlations between ROIs

We estimated pre-onset activity in V1 and the pulvinar (**Fig S4**), as well as conducting cross correlational analyses between given pairs of ROIs (**Figs 3D** **and S7**) in Matlab (2011b). For these analyses, we used the preprocessed raw time course data (see *fMRI processing*) which were z-normalised without deconvolution, as deconvolution would remove activity unrelated to stimulus onset which is the interest of analysis here. To estimate activity prior to the face onsets, the normalised signal change (%) in the time course was averaged between the 2 time points immediately before the face onsets (i.e., −5 and −2.5 s) relative to the preceding baseline (averaged between the 2 earlier time points, i.e., −10 and −7.5 s).

We further conducted time-shifted cross correlation analyses between V1 and the pulvinar (**Fig 3D**). Time course data for each area was time-shifted to examine which direction of the interactions (V1-to-pulvinar or pulvinar-to-V1) contributed to a false percept of a fearful face. This analysis was based on the assumption that the direction of interaction can be inferred when activity in one brain area shows greater correlation with the activity in another area from a certain past time point (33), as has previously been used to examine geniculocortical interactions (54). If pulvinar activity precedes V1 activity, the original (non-shifted) time course for V1 is expected to show greater correlation with the pulvinar time course that is shifted to an earlier time point. Conversely, if V1 activity precedes pulvinar activity, the original time course for the pulvinar is expected to show greater correlation with the V1 time course shifted to an earlier time point. Here, we used a basic correlation method rather than other related methods, such as Granger-causality analysis, as the latter are typically more optimal for datasets with higher temporal resolution (55).

To estimate correlation, we shifted the time courses of V1 and the pulvinar (originally synchronized with stimulus onsets) by a maximum or 1TR with an increment of 625 ms (i.e., 1/4 of TR), while linearly interpolating the data points in between TRs. The time lag was kept smaller than 1 TR to avoid potential overlap with non-sensory events such as key response. The data points in the time course covering the temporal peak of activity from the onsets of faces (2.5 to 7.5 s) were accumulated across trials for each combination of percept type (HIT/FA) and facial emotion (fearful/happy) to estimate the Fisher-transformed Pearson correlation coefficient (**Fig 3C**). Given that both the pulvinar and visual cortices typically respond to the contralateral visual fields (56, 57), the interaction between the pulvinar and V1 is expected to be confined within each hemisphere. Therefore, the Fisher-transformed correlation coefficient was estimated for each hemisphere, e.g., between the left pulvinar and left V1, and was subsequently averaged between the hemispheres. For participants who had a pulvinar or V1 ROI located in only one hemisphere (right only, N = 1; left only, N = 3), the cross correlation was estimated for that hemisphere. Cross correlations for a given pair of control ROIs were analysed in the same manner and are presented in **Fig S7**.

Although the difference in hemodynamic responses between the cortical and subcortical areas (58) may have affected the cross-correlation between V1 and pulvinar, such difference cannot explain away the enhanced correlation on FA trials relative to HIT trials of a fearful face.

## Author Contributions

AK, HB, BDG, and KA contributed to the study design, FDM and IK contributed to the scanning and parameter settings of 7T fMRI, AK and MZ collected the data, AK conducted the data analyses with guidance and feedback from FDM, MZ, HB, and KA. AK, KA, and BDG wrote the manuscript with feedback from HB, MZ, FDM, and IK.

## Acknowledgements

This research was funded by a European Research Council (ERC) grant under the European 512 Union Seventh Framework Programme for Research 2007-2013 (grant agreement number 513 295673), by European Union’s Horizon 2020 Research and Innovation Programme under grant 514 agreement No 645553, ICT DANCE (IA, 2015-2017), by Incentive grant of National Institute of Information and Communications Technology (NICT), Japan, and JSPS KAKENHI grant number 16K21687 and 18H02714 awarded to AK. The funders had no role in study design, data collection and analysis, decision to publish, or preparation of the manuscript. We thank Dr. Maarten Vaessen for his help in the pilot stage, and Prof. Yuka Sasaki, Prof. Ben Seymour, Prof. Kazuhisa Shibata, Prof. Nim Tottenham, Dr. Brian Maniscalco, and Dr. Jorge Morales for their useful comments.

